# Nonclinical Safety and Immunogenicity of an rVSV-ΔG-SARS-CoV-2-S vaccine in mice, hamsters, rabbits and pigs

**DOI:** 10.1101/2021.07.06.451119

**Authors:** Noa Madar-Balakirski, Amir Rosner, Sharon Melamed, Boaz Politi, Michal Steiner, Hadas Tamir, Yfat Yahalom-Ronen, Elad Bar David, Amir Ben-Shmuel, Assa Sittner, Itai Glinert, Shay Weiss, Erez Bar-Haim, Hila Cohen, Uri Elia, Hagit Achdout, Noam Erez, Shahar Rotem, Shlomi Lazar, Abraham Nyska, Shmuel Yitzhaki, Adi Beth-Din, Haim Levy, Nir Paran, Tomer Israely, Hadar Marcus

## Abstract

rVSV-ΔG-SARS-CoV-2-S is a clinical stage (Phase 2) replication competent recombinant vaccine against SARS-CoV-2. Nonclinical safety, immunogenicity and efficacy studies were conducted in 4 animal species, using multiple dose levels (up to 10^8^ PFU/animal) and various dosing regimens. There were no treatment related mortalities in any study, or any noticeable clinical signs. Compared to unvaccinated controls, hematology and biochemistry parameters were unremarkable and no adverse histopathological findings gave cause for safety concern in any of the studies. There was no viral shedding in urine, nor viral RNA detected in whole blood or serum samples 7 days post vaccination. The rVSV-ΔG-SARS-CoV-2-S vaccine immune response gave rise to neutralizing antibodies, cellular immune response, and increased lymphocytic cellularity in the spleen germinal centers and regional lymph node. No evidence for neurovirulence was found in C57BL/6 immune competent mice or in highly sensitive IFNAR KO mice. Vaccine virus replication and distribution in K18 hACE2 transgenic mice showed a gradual clearance from the vaccination site with no vaccine virus recovered from the lungs. The rVSV-ΔG-SARS-CoV-2-S vaccine was well tolerated locally and systemically and elicited an effective immunogenic response up to the highest dose tested, supporting further clinical development.

## 1. Introduction

Severe acute respiratory syndrome coronavirus 2 (SARS-CoV-2), a member of the Coronaviridae family, is responsible for the COVID-19 pandemic that emerged from the Wuhan Province in China in 2019 and rapidly spread globally, with over 175 million confirmed cases of COVID-19, including 3.8 million deaths, reported to WHO as of 14 June 2021, and a total of 2.3 x 10^9^ vaccine doses have been administered.

The urgent need for an efficacious vaccine for SARS-CoV-2 led to unprecedented innovation worldwide. Thirteen vaccines have been approved under emergency use authorizations (EUA) in different countries [1]. The U.S. Food and Drug Administration has granted EUA for three COVID-19 vaccines so far; Pfizer-BioNTech’s vaccine (December 11, 2020); Moderna’s vaccine (December 18, 2020) and Janssen’s vaccine (February 27, 2021); Astra Zeneca received conditional marketing approval for its vaccine throughout the European Union (January 29 2021).

However, while several countries—including Israel, the United Kingdom, the and Canada, have made significant progress towards vaccinating a large portion of their citizens (with over 60% of their population that received at least one vaccine dose) [2], for most of the countries of the world, vaccines are yet not available. It is, therefore, essential to develop vaccines, to meet this unmet need in a timely manner. As of May 9th 2021, nearly 300 vaccines were in development using a variety of technologies, including over 100 vaccines that are currently tested in clinical trials [3].

We recently presented the development of rVSV-ΔG-SARS-COV-2-S vaccine, currently in phase 2 clinical trials in Israel. rVSV-ΔG-SARS-COV-2-S is a vector-based vaccine that utilizes the recombinant vesicular stomatitis virus (VSV) platform in which the glycoprotein gene of VSV (VSV-G) was genetically replaced by the human codon optimized spike protein gene of SARS-CoV-2 [4]. The approach for developing this vaccine is similar to the one used by Merck Sharp & Dohme B.V. for Ervebo®, a vaccine developed against Ebola virus, in which the VSV-G was replaced by that of Zaire Ebola Virus Kikwit 1995 strain glycoprotein [5]. The efficacy of the vaccine was demonstrated using the golden Syrian hamster in-vivo model for COVID-19, where it was shown that a single-dose vaccination with the rVSV-ΔG-SARS-CoV-2-S vaccine resulted in a rapid and potent induction of SARS-CoV-2 neutralizing antibodies. In addition, vaccination protected hamsters against SARS-CoV-2 challenge, as demonstrated by the abrogation of body weight loss in immunized, compared to unvaccinated hamsters, and alleviation of the extensive tissue damage and viral loads in lungs and nasal turbinates [6].

In order to evaluate the safety profile of the rVSV-ΔG-SARS-COV-2-S vaccine, and to support its clinical development and licensure, a series of nonclinical safety, immunogenicity and efficacy (potency) studies were performed. In this work, we present the results of the nonclinical safety and immunogenicity studies conducted in 4 animal species; mice, hamsters, rabbits and pigs. The design of the studies was based on the principles described in the WHO Guideline of Nonclinical Evaluation of Vaccines (Annex 1) [7], which is recognized by both FDA and EMA as well as by the EMA Guideline on quality, non-clinical and clinical aspects of live recombinant viral vectored vaccines [8], and included single and repeated dose studies. In addition, attention was given to potential risks that may be associated with viral vectored vaccines, including the potential of neurovirulence, assessment of vaccine shedding, vaccine replication and potential distribution to tissues of interest.

## 2. Materials and Methods

### 2.1 Virus and vaccine

Generation of the pVSVΔG-S construct was described in detail, previously [6]. Briefly, the pVSV-spike expression plasmid was constructed by PCR amplification of the full-length human codon-optimized S gene from pCMV3-SARS-CoV-2 S expression plasmid (Sino Biological, Cat# VG40588-UT) that was used to replace the VSV-G (Glycoprotein) open reading frame within the VSV full length expression vector, yielding pVSV-ΔG-spike. Primary recovery of the rVSV-ΔG-SARS-CoV-2-S was performed in BHK-21 cells infected with Modified Vaccinia Ankara T7 (MVA-T7), followed by co-transfection with the rVSV-SARS-CoV-2-S, and the VSV accessory plasmids encoding for VSV-N, P, L and G proteins under control of a T7 promoter.

### 2.2 Animals

C57BL/6J, knockout for type I interferon (IFN) receptors (Ifnar−/−) and hACE2-K18 mice were obtained from The Jackson Laboratory, USA, and were 6-12 weeks of age at study initiation. Golden Syrian hamsters were obtained from Charles River, USA, and were 6-7 weeks of age at study initiation.

Female New Zealand White (NZW) rabbits were obtained from Charles River, France, and were 14-16 weeks of age at study initiation. Female pigs (Topigs 20, a cross between the Z line, Large-White dam and the N-line, Landrace) were obtained from Van Beek, Netherlands, and were 9-10 weeks of age at study initiation.

Animals were housed in group cages in a biosafety level 2 (BSL-2) containment facility, maintained at room temperature between 18 to 24°C and 30 to 70% relative humidity, with a 12-h light/12-h dark cycle, and received a pelleted diet. Mice, hamsters and rabbits had free access to food and water throughout all the experiments. Pigs were fed twice a day (4% of their body weight daily) and allowed free access to drinking water supplied by automated watering valves.

### 2.3 Ethics statement

Experiments were approved by the IIBR animal care and use committee (IACUC) and performed in accordance with the guidelines of the care and use of laboratory animals published by the Israeli Ministry of Health (protocols: mice C57BL/6J # No. M-52-20, 3 September 2020; Ifnar1 knock-out and hACE2-K18 mice # No. M-65-20, 12 October 2020; hamster # No. HM-02-20, 6 April 2020; rabbits # No. RB-13-29, 28 May 2020; pigs # No. P-03-20, 4 May 2020).

### 2.4 Procedures and Examinations

Animal Vaccination: Anesthetized golden Syrian hamsters were vaccinated by a single i.m. injection (hind shin) at a dose of 10^6^ PFU and at a dose volume of 0.05 ml/animal. Pigs and rabbits were vaccinated by 2 successive i.m. injections (thigh muscles and shin area, respectively) at a dosing interval of 2 weeks, at dose levels of 10^6^ – 10^8^ PFU (see details in the results section) and at a dose volume of 1ml/animal (single injection site). K18 hACE2 transgenic mice were vaccinated by a single i.m. injection (hind limb) at a dose of 10^7^ PFU and at a dose volume of 0.05 ml/animal. Body weight, general observations and local reactions at injection sites were monitored throughout each study period. Body temperature was also measured in the rabbits and pigs.

Control animals were injected with carrier buffer at an identical dose volume and under identical experimental conditions.

### 2.5 Immunogenicity and Cellular Immunity Determination

The immunogenicity of the vaccine was evaluated by determining the NT50 values of neutralizing antibodies against SARS-CoV-2 (the dilution at which 50% neutralization was observed), as was assessed by the plaque reduction neutralization test (PRNT) [6]. Cellular immunity was analyzed in vaccinated and naïve pigs. Peripheral blood mononuclear cells (PBMC) were stimulated for 48 hours in the presence of spike protein (0, 1 or 10 μg) [9]. Interferon γ (IFNγ)-secreting cells were enumerated with Pig IFN-γ Single-Color ELISPOT kit (ImmunoSpot, CTL Technologies, Germany) with strict adherence to the manufacturer’s instructions. The frequency of cytokine-secreting cells was quantified with ImmunoSpot S6 reader and analyzed with the ImmunoSpot software (ImmunoSpot, Cellular Technology Limited, Germany). Secreted cytokine (IL-2, IFNγ, IL-4, IL-10 concentrations were determined with MILLIPLEX MAP Porcine Cytokine/Chemokine Magnetic Bead Panel (Merck-Millipore) with strict adherence to the manufacturer’s instructions.

### 2.6 Hematology, Biochemistry, Coagulation, Viremia and Viruria

Blood was collected from rabbits and pigs via the central (auricular) artery or the marginal ear veins, after applying local anesthesia to the outer ear (e.g. Emla 2.5 % Topical Cream, applied 30 minutes prior to blood sampling) and following pre-heating of the animal environment (for about 5 minutes). Blood collected for hematology was discharged into an EDTA tube (inverted and stored on wet ice) blood collected for biochemistry was discharged into serum separator tube. The serum was allowed to clot at room temperature for at least half an hour; blood was centrifuged at 1000 g for 15 minutes and serum poured off into 2 labeled 1.2 mL cryogenic vials, immediately analyzed or kept and frozen at or below −70°C for future analysis (viremia).

For hematological analysis, a complete blood count, including platelet and differential count was performed on the day of collection (Oxford Science GenX, USA). For clinical biochemistry, serum samples were evaluated for the following endpoints: albumin, alkaline phosphatase alanine aminotransferase, amylase, total bilirubin, blood urea nitrogen, calcium, phosphorous, creatinine, glucose, sodium, potassium, total protein and globulin. For coagulation, serum samples were evaluated for the following endpoints: prothrombin time (PT), partial prothrombin time (PTT) and fibrinogen (Abaxis- Vet Scan VS2, USA).

Whole blood and urine samples were collected for assessment of viremia and viruria, respectively, at several timepoints (see results section). Whole blood was collected using the same procedures as described above, and urine samples were collected from the cage’s tray. Samples were prepared as follows: 200 μl of urine or blood sample were added to 150 μl of lysis buffer (LBF, supplied with the extraction kit) and incubated for 20 minutes at room temperature to complete lysis. Following lysis, RNA extraction was completed using an RNAdvance Viral kit as per manufacturer’s instructions. Real time RT-PCR was performed using the SensiFAST™ Probe Lo-ROX one-step kit. In each reaction the primers final concentration was 600 nM and the probe concentration was 300 nM. Primers and probes were designed using the Primer Express Software (Applied Biosystems).

### 2.7 The primers and probes used

N gene: forward: TGATCGACTTTGGATTGTCTTCTAA, reverse: TCTGGTGGATCTGAGCAGAAGAG, probe: ATATTCTTCCGTCAAAAACCCTGCCTTCCA; S gene: forward: GAGTGAGTGTGTGCTGGGACAA, reverse: AAACACTCCCTCCCTTGGAAA, probe: AGTTTTCCACAGTCTGCCCCTCATGGA.

The probes used were 6-FAM and ZEN/Iowa Black FQ combination. Assay performance was verified using appropriate controls.

### 2.8 Macroscopic and histopathological examinations

Hamsters were euthanized and the following organs: brain, heart, lung, liver, kidney, colon and the shin-injection site were collected. Organs were fixed in 4% formaldehyde, trimmed in a standard position per organ and placed in embedding cassettes. Paraffin blocks were sectioned at approximately 4 microns thickness. The sections were placed on glass slides and stained with Hematoxylin & Eosin (H&E). Histopathological evaluation of the slides was performed by PATHO-LOGICA Ltd.

Rabbits and pigs were subjected to a full detailed necropsy and gross pathological examination following the respective scheduled termination. At necropsy, all animals were thoroughly examined, including the external surface of the body, all orifices, cranial, thoracic and abdominal cavities and their contents. Any abnormalities or gross pathological changes observed in tissues and/or organs were recorded, accordingly. All organs/tissues were collected from all animals during necropsy and fixed in 10% neutral buffered formalin (approximately 4% formaldehyde solution) apart from the eyes, which were fixed in Davidson’s Solution. All organs/tissues were trimmed, embedded in paraffin, sectioned at approximately 5 microns thickness and stained with Hematoxylin & Eosin (Alizée Pathology, Inc). The slides were subjected to histopathological evaluation by a Board Certified toxicologic pathologist, using the state of the art recommended morphological criteria published by the International Harmonization of Nomenclature and Diagnostic Criteria (INHAND), https://www.toxpath.org/inhand.asp. A grading scheme, derived from Schafer et al., 2018 [10], was used to evaluate pathologic lesions in the tissues as follows: no lesion (0), minimal (grade 1), mild (grade 2), moderate (grade 3) and marked (grade 4).

### 2.9 Vaccine Replication in the K18 hACE2 transgenic mouse model

Animals were euthanized by cervical dislocation immediately following vaccination (time 0), 24, 48 and 72 hours post vaccination (n=4 mice per time point). The site of injection and the lungs were excised, then snap frozen (liquid nitrogen followed by storage at −70°C). The organs were processed for titration in 1.5ml ice-cold PBS and titered on VERO-E6 cells. PFUs were calculated according to the sample dilution as described previously [11].

### 2.10 Neurovirulence testing

VSV-WT or rVSV-ΔG-SARS-CoV-2-S vaccine were injected intracranially to C57BL/6 immune competent mice (6-12 weeks old, n=3/dose level) and to type I interferon receptor knock-out (IFNAR KO) mice (6-12 weeks old, n=3/dose level) at a dosing volume of 30μl. Naïve mice, and mice which were injected with the formulation buffer only (n=3/group for each mouse strain) were used as controls. Mice were monitored daily for safety and morbidity for 14 consecutive days. For histopathology evaluations, rVSV-ΔG-SARS-CoV-2-S C57BL/6 (n=3), rVSV-ΔG-SARS-CoV-2-S IFNAR (n=6) or VSV-WT C57BL/6 (n=5) intracranially injected mice were sacrificed on day 14, 14 and 2, respectively. In addition, vehicle-injected (n=3/strain) and naïve (n=3/strain) mice were sacrificed on day 14 and served as controls. Mice were sedated, perfused and fixed with 4% formaldehyde, and brains were collected and fixed with 4% formaldehyde for histopathological examinations of appropriate areas. Paraffin blocks were transversely sectioned at approximately 4 microns thickness. For each brain, three transverse sections were cut at the level of the striatum (Sr), hippocampus (Hc) and cerebellum (C), to include the frontal cortex (FC), striatum (Sr), thalamus (TH), hypothalamus (H), hippocampus (Hc) and cerebellum (C). Sections were placed on glass slides and stained with Hematoxylin & Eosin (H&E) for general histopathology. The slides were evaluated by the IIBR histology unit. H&E stained sections were examined and graded according to a semi-quantitative scoring scale for the presence and severity of pathological changes. Histopathologic changes consistent with neurovirulence were scored, in a double blinded manner, based on the presence and severity of perivascular infiltrates, gliosis, neurodegeneration, satellitosis, and necrosis. Cumulative assessment of all changes was scored on a severity scale of 0– 4 based on previously published work [12].

## 3. Results

### 3.1. rVSV-ΔG-spike vaccine safety in a COVID-19 hamster model

We recently reported that a single-dose vaccination with rVSV-ΔG-SARS-CoV-2-S vaccine resulted in a rapid and potent induction of SARS-CoV-2 neutralizing antibodies in a golden Syrian hamster model, and protected hamsters against SARS-CoV-2 challenge, as demonstrated by the abrogation of body weight loss, alleviation of the extensive tissue damage and viral loads in lungs and nasal turbinates [6]. We further utilized this model in order to evaluate the safety of the candidate vaccine, in three groups of female Syrian hamsters. One group of hamsters (n=20 females) was administered with rVSV-ΔG-SARS-CoV-2-S vaccine by a single i.m. injection at a dose of 106 PFU and at a dose volume of 0.05 ml/animal. A vehicle control group (n=4 females) was administered with the carrier buffer (PBF solution) and a naïve control group (n=4 females) served as an additional control group. The dose of 106 PFU was selected as it was shown to be immunogenic and protective in this model [6]. Following administration of rVSV-ΔG-SARS-CoV-2-S vaccine, no treatment related mortality, nor noticeable systemic or local reactions were noted in any of the animals throughout the 24-day observation period. The vaccinated animals exhibited mean group body weight gain similar to that of the control groups, 7 days post vaccination (Figure 1A). Histopathological evaluation carried out 7 days post vaccination revealed no treatment-related pathological, cytotoxic or other adverse effects in the tested organs (brain, heart, lung, liver, kidney, colon and the shin-injection site) in all treated animals. Blood samples were obtained from 5 vaccinated hamsters, 20 days post vaccination for determination of neutralizing antibody titer. Neutralizing antibodies were detected in all vaccinated hamsters, although high variability was observed (n=5, Mean=2371, SEM=806.6 Range: 301.9-5133. Figure 1B). Taken together, a single-dose vaccination of hamsters with rVSV-ΔG-SARS-CoV-2-S vaccine (106 PFU/animal) was found to be safe and immunogenic and did not lead to any treatment-related histopathological changes in the main organs tested (data not shown).

**Figure 1.**
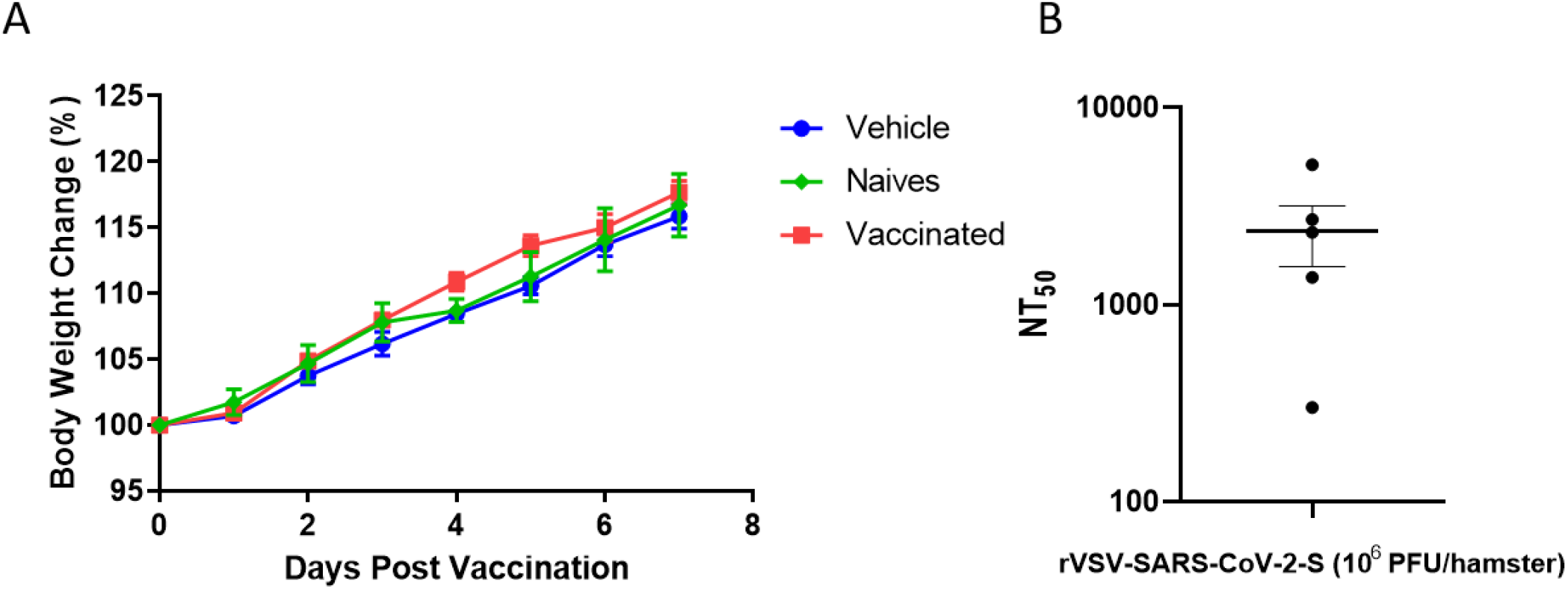
Vaccinated hamsters body weight changes and serum neutralizing antibody titers. (a) Body weight changes of vaccinated hamsters (10^6^ PFU, n=20) compared to control animals (naive or mock-vaccinated, n=4/group) throughout the first 7 days post vaccination. (b) NT50 values of i.m. vaccinated hamsters’ sera, data presented as mean values ± SEM (10^6^ pfu/hamster, n = 5).

### 3.2. Safety and Immunogenicity in NZW Rabbits

Four groups of female NZW rabbits (n= 4 females/group) were tested. Three groups were vaccinated with rVSV-ΔG-SARS-CoV-2-S vaccine at dose levels of 10^6^, 10^7^ or 10^8^ PFU (1 ml/animal/injection). An additional group was injected with the vehicle and served as control. All animals were subjected to 2 repeated i.m. vaccinations (prime and boost) at an interval of 2 weeks, and follow-up continued for an additional 21-day recovery period (total observation period of 35 days). Body weight, general observations, local reactions at injection sites and body temperature were monitored throughout the study period. No mortality occurred and no noticeable systemic or local reactions were noted in any of the animals throughout the 35-day study period. All rVSV-ΔG-SARS-CoV-2-S vaccine treated groups exhibited body weight gain comparable to that of the control group (Figure 2A), and mean group body temperature values of all groups were within the normal range of body temperature throughout the seven days post prime and boost vaccinations. (Figure 2B) [13].

**Figure 2.**
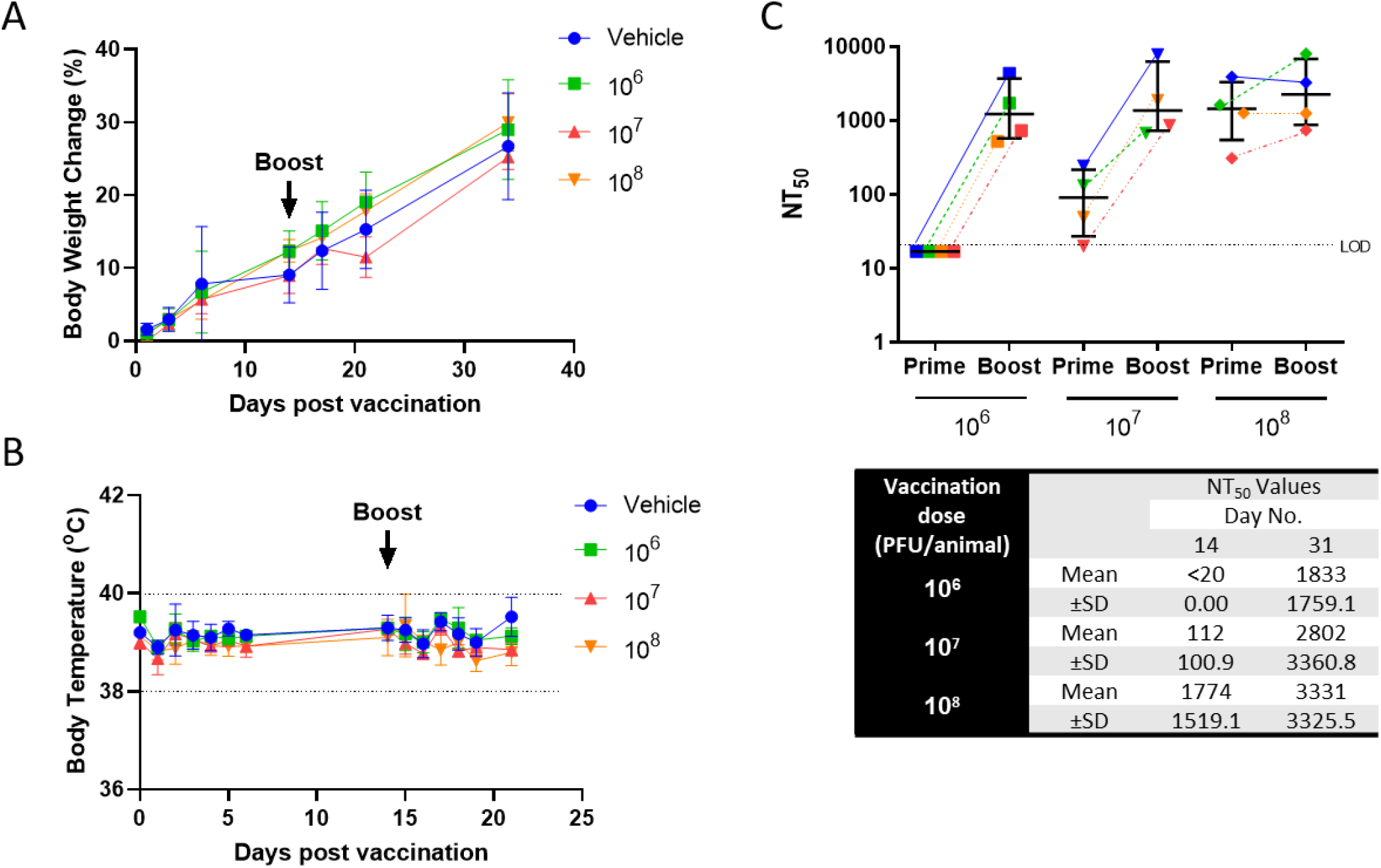
Body weight, temperature and serum neutralizing antibody titers in rabbits after prime/boost vaccination. (a) Body weight change percentage compared between groups (n=4 /group). (b) Group mean body temperature (°C) following prime and boost vaccination of rabbits. Dashed lines indicate normal body temperature range. (c) vaccinated rabbit sera neutralization assay results [mean group NT50, using plaque reduction neutralization test (PRNT)]. Sera were collected on day 14 (14 days post first vaccination; “prime”) and on day 31 (17 days post the second vaccination; “boost”). Dotted line represents the limit of detection (LOD). Error bars show median interquartile range (IQR). Means and SD are indicated below the graph.

Serum samples were collected on day 14 (prior to vaccination boost) and on day 31, 17 days after the vaccination boost and were analyzed for the presence of neutralizing antibodies against SARS-CoV-2. Dose– response dependency in mean group neutralizing antibodies was evident following each vaccination session and mean group NT50 was increased in response to the boost vaccination in the low and medium dose groups. Following vaccination with the high dose (10^8^ PFU), neutralizing titers were efficiently induced following prime vaccination, and boost vaccination did not significantly affect the neutralization titer (Figure 2C).

Hematology, biochemistry and coagulation parameters were assessed at several timepoints. In all vaccine treated groups these parameters were found to be similar to the baseline or control animals’ values throughout the study period. The potential for viral shedding in urine samples (viruria) and in whole blood and serum samples (viremia) was tested in order to evaluate the extent of the vaccine dissemination from the site of administration, the rapidity of its clearance and its potential dissemination through secretions. No evidence of viremia was noted in any of the serum samples obtained 3 and 7 days post the prime vaccination, in addition, no shedding of viral vaccine was detected on day 3 post the prime vaccination in any of urine samples collected (data not shown).

At necropsy, 21-day post second vaccination, no treatment related macroscopical findings or changes in organ weight and organ to body weight ratio values were evident in any of the vaccine treated animals. Histopathological evaluation revealed treatment-related changes in all treated groups, generally being dose-related in incidence and/or severity, and which corresponded to the pharmacological activity of the vaccine. The changes were seen at the injection sites, regional iliac lymph nodes and spleen (Figure 3).

**Figure 3.**
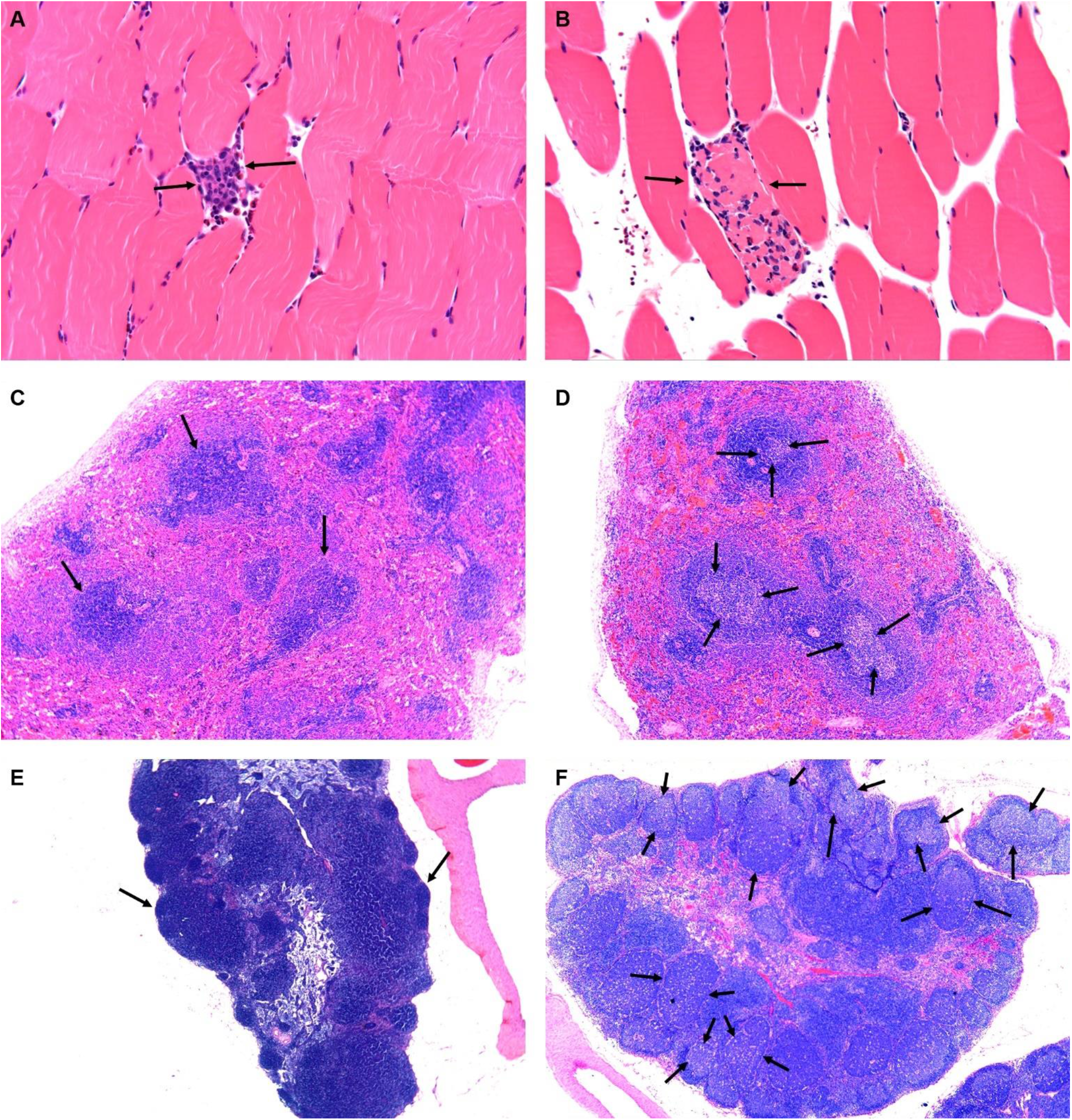
Rabbit histopathology following prime/boost vaccination. Histopathological evaluation performed 3-weeks post second vaccination session. (a-b) Injection Site Analysis (high dose, 10^8^ PFU/animal)-arrows indicate changes at the injection sites (shin area) consisting of focal minimal mixed inflammatory cell infiltration (a) and focal minimal muscle fiber necrosis associated with mixed inflammatory cell infiltration (b) which are not considered as adverse treatment effects. (c) Spleen of control animal and (d) high dose (10^8^ PFU) animal: arrows indicate minimal to mild germinal centers increased lymphocytic cellularity (i.e. follicular hyperplasia), noted following vaccination and considered to reflect a secondary change due to antigenic stimulation. No evidence of germinal centers were observed in a control animal (c). (e) Iliac Lymph Node (regional lymph node to the injection site) of control animal and (f) of the high dose (10^8^ PFU) animal: arrows indicate mild to moderate germinal centers increased lymphocytic cellularity (i.e. follicular hyperplasia), noted following vaccination and considered to reflect a secondary change due to antigenic stimulation. No evidence of germinal centers were observed in a control animal (e).

The changes at the injection sites consisted of multifocal minimal myofiber necrosis (limited to the 2 higher doses); multifocal minimal to mild mixed inflammatory cell infiltration (i.e. cells consisting of a mixture of polymorphonuclear cells, lymphocytes and macrophages within the skeletal muscle and/or interstitial fat tissue) and focal minimal granulomatous inflammation (seen in a single Low-dose and in a single High-dose animal). This inflammatory reaction was also seen in the fat tissue surrounding the sciatic nerve, but did not extend into the sciatic nerve itself. This change is considered an extension of the primary injection site inflammation, possibly related to the needle location/insertion and is not considered to reflect any increase in local irritation (Figure 3 A-B).

In the spleen, minimal to mild germinal centers increased lymphocytic cellularity (i.e. follicular hyperplasia) were noted (Figure 3 C-D). In the iliac lymph nodes (i.e. regional lymph node to the injection site), mild to moderate germinal centers increased lymphocytic cellularity (i.e. follicular hyperplasia) were seen in all treated groups (Figure 3 E-F). The germinal centers increased lymphocytic cellularity (i.e. follicular hyperplasia), seen in spleen and iliac lymph nodes, is considered to reflect a secondary change due to antigenic stimulation of the test compound [14], and were reported previously in rabbits exposed to vaccination [15].

### 3.3. Safety and Immunogenicity in Domestic Pigs

In this study, 2 groups of female Topig 20 pigs (n= 2 females/group) were vaccinated with rVSV- ΔG-SARS-CoV-2-S vaccine at dose levels of 10^6^ and 10^8^ PFU (1ml/animal, in a single injection site). An additional group (n= 2 females) was injected with the carrier buffer and served as control. Similar to the rabbit study, the pigs were subjected to 2 repeated i.m. vaccinations (prime and boost) at an interval of 2 weeks, and follow-up continued for an additional 18-day recovery period (total observation period of 32 days). No mortality occurred and no noticeable systemic or local reactions were noted in any of the animals throughout the 32-day study period. All rVSV-ΔG-SARS-CoV-2-S vaccine treated groups exhibited body weight gain comparable to that of the control group (Figure 4A), and the temperature profile of all animals was within the normal range [16] (Figure 4B).

**Figure 4.**
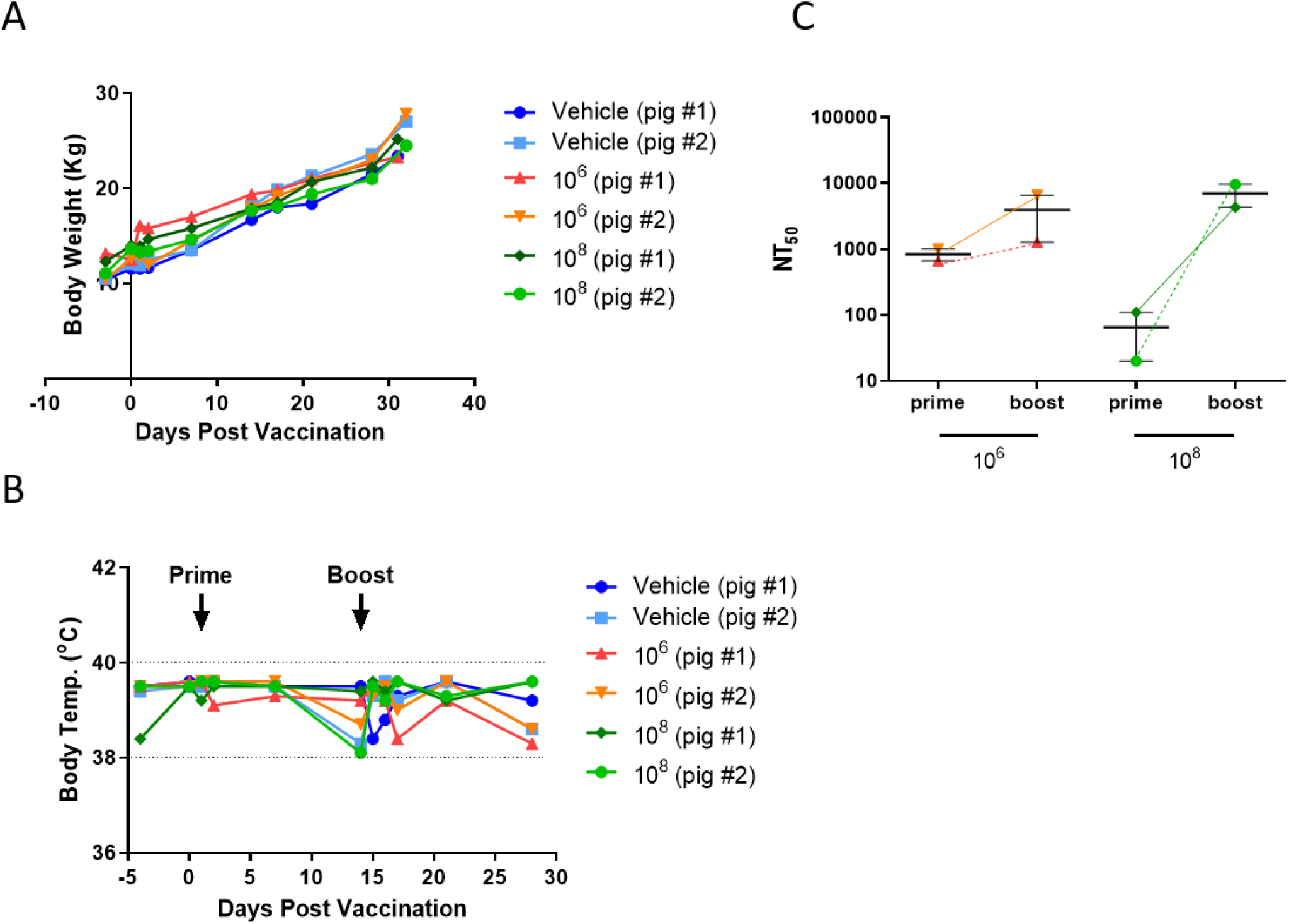
Body weight, temperature and serum neutralizing antibody titers in pigs after prime/boost vaccination. (a) Individual body weight change (percentage). (b) Individual body temperature (°C) following prime and boost vaccination of pigs. Dashed lines indicate normal body temperature range. (c) Individual pig sera neutralization assay results (mean group NT50, using plaque reduction neutralization test (PRNT). Sera were collected on day 14 (14 days post first vaccination; “prime”) and on day 31 (“boost”). Error bars show median interquartile range (IQR).

Blood samples for immunogenicity were collected on day 14 (prior to vaccination boost) and on day 31, 17 days after the vaccination boost, and were analyzed for the presence of neutralizing antibodies and for cellular immunity responses.

A positive association was found between the neutralization antibodies levels and the stimulation of the cellular immunity. Both rVSV-ΔG-SARS-CoV-2-S vaccine doses induced a neutralization response (Figure 4C) and an increase in T-cell response to spike protein (Figure 5). Significant levels of IFNγ-secreting cells were observed among all the vaccinated pigs, up to 4×10^3^ cells per 10^6^ PBMC, following boost. The boost effect was observed in all the vaccinated animals, in accordance with increased neutralization antibodies levels (Figure 5A). Interleukin-2 and TNFα secretion was evident following spike antigenic stimulation in all the vaccinated pigs, with no IL-4 and marginal IL-10 secretion, indicative of a clear Th-1 response, (Figure 5B). Hematology, biochemistry and coagulation parameters were assessed at several timepoints throughout the study and were generally similar to baseline values or to control animals’ values. There was no evidence of viremia in either whole blood (day 2 and 31) or serum (day 2, 7, 14 and 31), or viral shedding in urine samples (day 2/3 and 7/8) throughout the study period.

**Figure 5.**
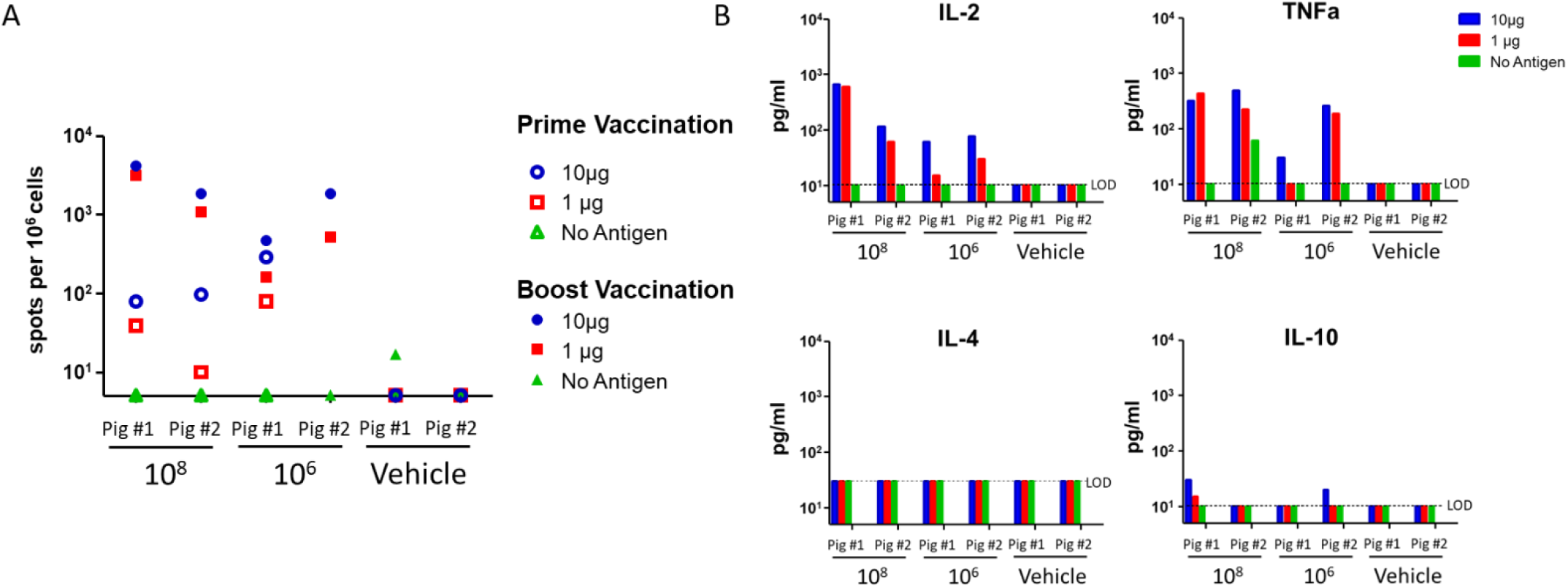
Cytokine secretions in pigs after prime/boost vaccination. (a) IFNγ secretion with Elispot assay: Antigen induced IFNγ producing T cells (spots/10^6^ PBMCs). (b) Cytokine secretion 3 weeks post-booster dose in pigs: Cytokine measurements were in duplicates for no antigen controls and naïve animals, and were in quadruplates for pigs vaccinated with rVSV-SARS-CoV-2-S. Positive controls (not shown) were measured for each cytokine in duplicate.

At necropsy and histopathology evaluation, no treatment related findings were observed. Sporadic minimal spontaneous lesions were noted in the liver, i.e. minimal inflammatory cell infiltration. Such a change is reported to occur spontaneously in pig/minipigs. In the spleen and lungs congestion was noted, which was comparable in control and treated animals, and is considered as related to the termination technique, injection of barbiturates, and not related to treatment. Such lesions are very well known to be associated with barbiturate injection for euthanasia [17].

### 3.4. Replication of rVSV-ΔG-SARS-CoV-2-S vaccine in K18 hACE2 transgenic mice

To further evaluate the safety of rVSV-ΔG-SARS-CoV-2-S vaccine, the ability of the vaccine to propagate at the site of injection and to spread to the lungs was assessed in K18-hACE2 transgenic mice in which the human ACE-2 receptor (hACE2) is expressed under the K18 promotor, in all epithelial cells. K18-hACE2 transgenic mice are considered to be a highly sensitive animal model for SARS-CoV-2 [18–20], and were also shown to induce neutralizing antibodies in response to rVSV-ΔG-SARS-CoV-2-S vaccination (manuscript in preparation). Mice were vaccinated i.m. in the right shin limb with 10^7^ PFU in a volume of 50μl, and were euthanized immediately following vaccination (time 0), 24, 48 and 72 hours post vaccination (n=4 mice per time point). The site of injection and the lungs were collected and processed, and infectious virus was quantified by plaque assay, as described above [6].

A gradual decrease in the viral titer was observed at the site of injection (Figure 6). Out of the injected 10^7^ PFU, an average of 4.1×10^6^ (41%) were recovered immediately after injection. This reduction can be attributed to the technical procedures performed on all organs following retrieval. The viral load was then gradually reduced, until a value lower by more than 4 logs within 72 hours post infection was reached (0.003% of the dose determined at time zero). In lungs, virus was not detected at any time point, in all mice. These results demonstrated the existence of the vaccine at the site of injection and its gradual clearance (over 3 days) but also show that there was still viable virus up to 72 hours post vaccination in some of the animals – a time window which allows for formation and induction of a robust immune response.

**Figure 6.**
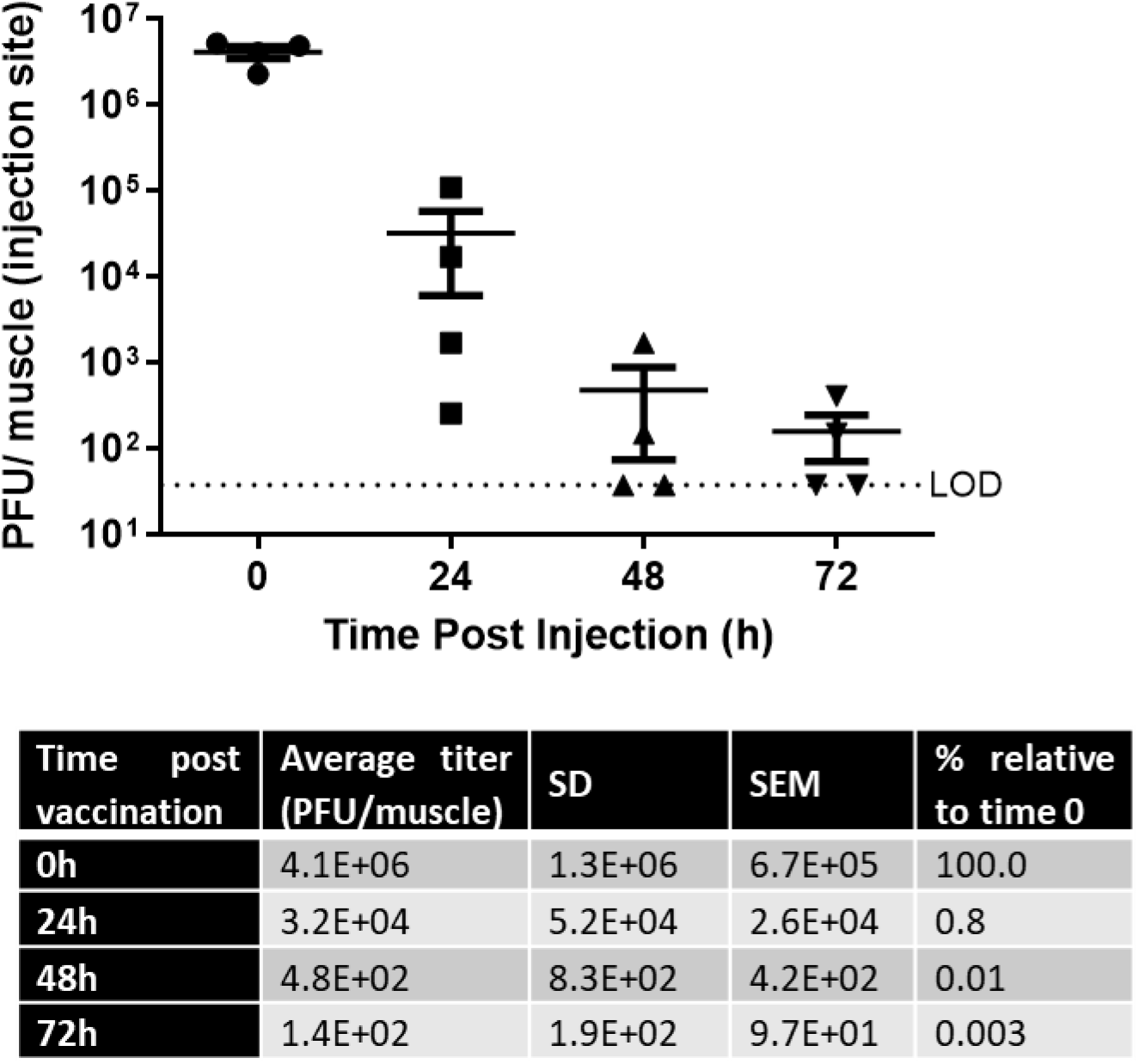
Vaccine viral load at site of injection in K18-hACE2 transgenic mice. Vaccine viral load was determined at in the injection site (muscle) at the time points indicated following vaccine administration to K18-hACE2 transgenic mice.

### 3.5. Neurovirulence in Mice

Several studies in cultured cells and mice demonstrated the role of VSV-G as a major virulent component of wild-type VSV, mediating infection of neurons and neurovirulence in mice. Consequently, when VSV-G was eliminated and replaced with a heterologous viral surface protein, neurovirulence was abrogated [21,22]. In ERVEBO®, elimination of VSV-G and replacing it with the glycoprotein (GP) of Ebolavirus (rVSV-ZEBOV-GP) rendered the viruses non-virulent and safe in general and especially safe to the nervous system as demonstrated by intra-thalamic injection in non-human primates, whereas wild-type VSV (VSV-WT) was highly neurovirulent [23]. To specifically pinpoint the safety of rVSV-ΔG-SARS-CoV-2-S vaccine in the nervous system, a comparative neurovirulence study was conducted by a single intracranial (i.c.) injection at various doses of either the wild-type VSV (VSV-WT) or rVSV-ΔG-SARS-CoV-2-S vaccine to C57BL/6 immune competent mice, and type I interferon receptor knock-out (IFNAR KO) mice. The latter are incapable of transducing signals through the interferon receptors alpha and beta and, as such, are highly sensitive to viral infections [24]

In C57BL/6 immune competent mice, a single intracranial (i.c.) injection of VSV-WT at doses of 10^3^, 10^4^ and 10^5^ PFU, resulted in rapid weight loss and death within 72 hours, while injection of a high dose of 105 PFU of rVSV-ΔG-SARS-CoV-2-S, was well tolerated, all mice survived and no reduction in body weight was noted (n=3/group; Figure 7A). In IFNAR KO mice, VSV-WT was uniformly lethal following a single intracranial (i.c.) injection at doses of 10^2^ or 10^3^ PFU/mouse. Mice rapidly deteriorated and died within 48-72 hours. IFNAR KO mice injected i.c. with rVSV-ΔG-SARS-CoV-2-S vaccine at doses of 10^3^ or 10^4^ PFU/mouse survived throughout the follow-up period. Mice did not show any signs of morbidity and no reduction in body weight was noted (n=3/group; Figure 7B).

**Figure 7.**
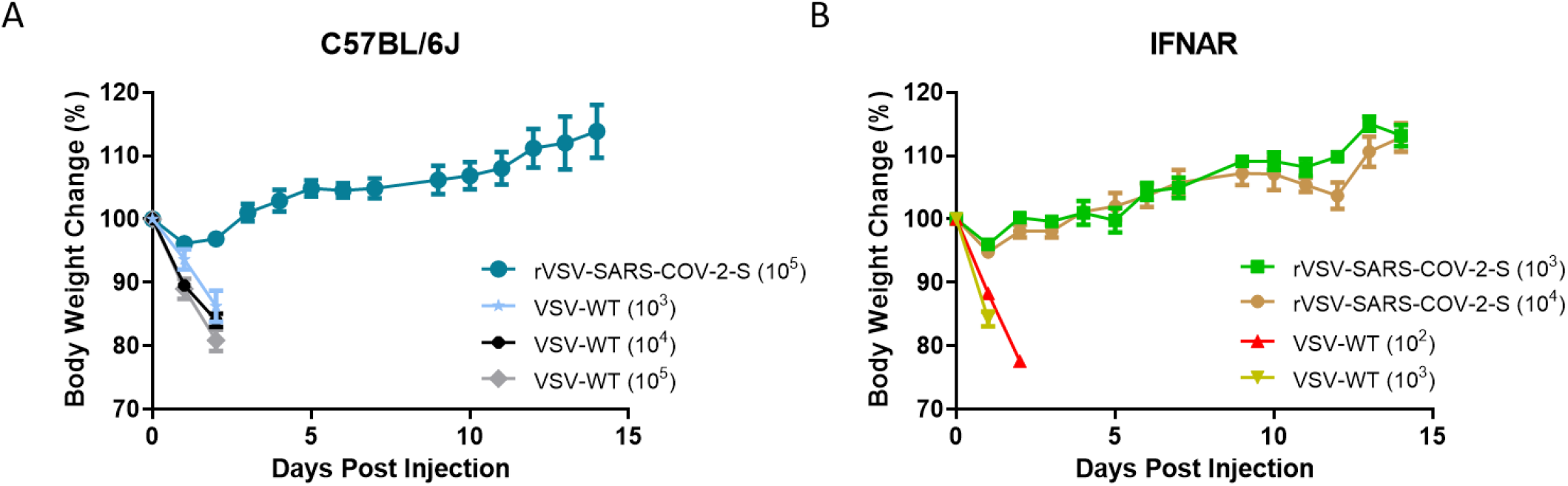
Comparison of VSV-WT with rVSV-ΔG-SARS-CoV-2-S vaccine effects following i.c. administration to C57BL/6J or IFNAR KO mice. (a) C57BL/6J mice, 6-12 weeks of age, administered i.c. with VSV-WT or rVSV-ΔG-SARS-CoV-2-S vaccine virus at doses ranging from 10^3^ to 10^5^ PFU (n=3/group). (b) IFNAR KO mice, 6-12 weeks of age, administered i.c. with VSV-WT or rVSV-ΔG-SARS-CoV-2-S vaccine virus doses ranging from 10^2^ to 10^4^ PFU.

Histopathology evaluation of the brains of both mice strains that were injected with rVSV-ΔG-SARS-CoV-2-S vaccine revealed no perivascular inflammatory infiltrate, no gliosis, no neurodegeneration, no satellitosis, and no necrosis in all the evaluated brain areas (including the frontal cortex, striatum, thalamus, hypothalamus, hippocampus and cerebellum); representative images from C57/BL6 injected mice are shown in Figure 8. Neurovirulence was absent in both C57BL/6 immune competent mice and IFNAR KO mice, injected with the rVSV-ΔG-SARS-CoV-2-S vaccine.

**Figure 8.**
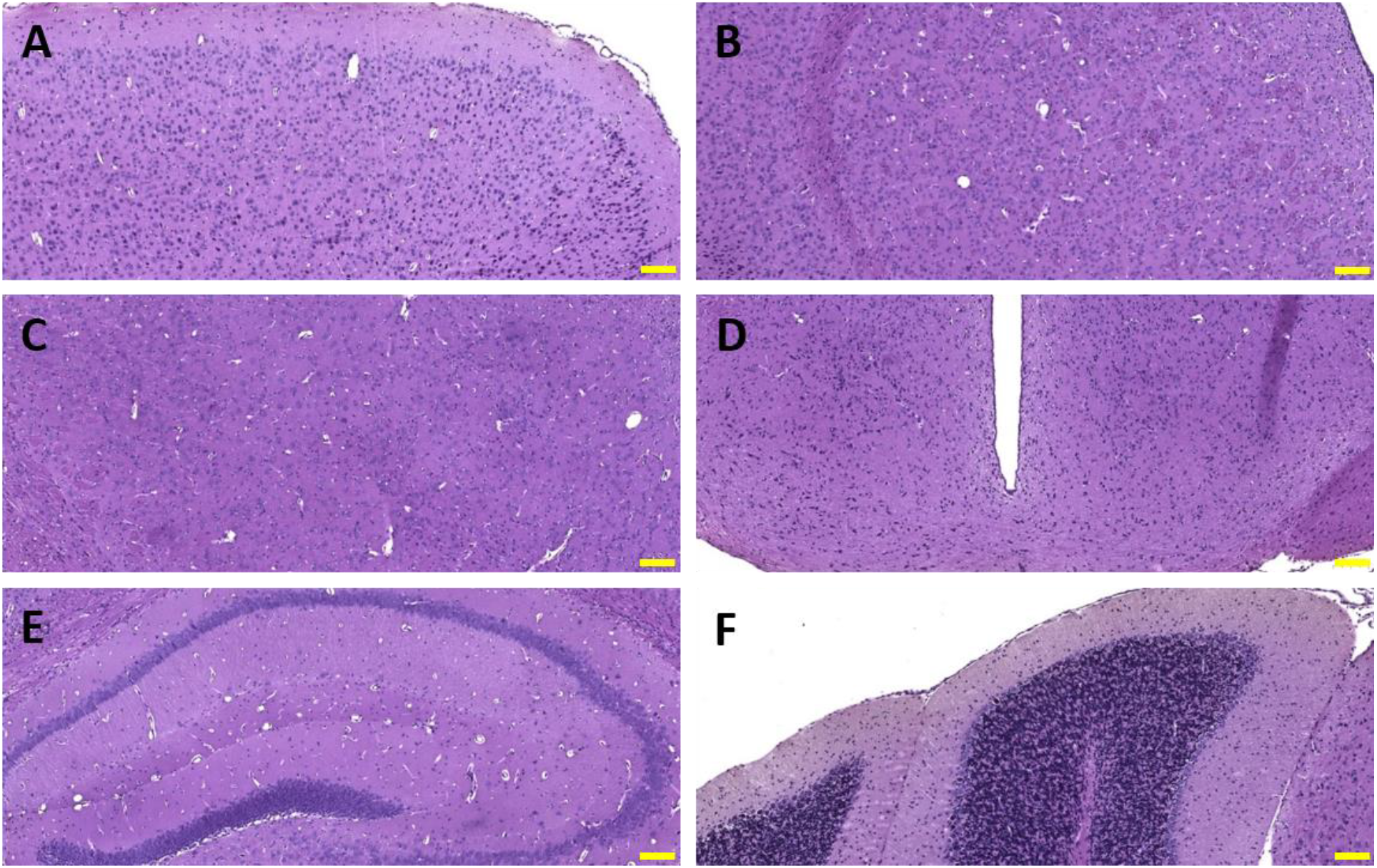
Histopathology of C57/BL6 mouse brain after injection with rVSV-ΔG-SARS-CoV-2-S. (a) Frontal cortex FC; (b) Striatum, Sr; (c) Thalamus, TH; (d) Hypothalamus, H; (e) Hippocampus, Hc; (f) Cerebellum, C. Scale bar=100μm, Magnification: X10, H&E staining.

In contrast, brains of mice injected with VSV-WT, presented with profound histological damage. Pathology was pronounced in the striatum (Sr) area, mainly in the Corpus Callosum (CC) and in the Medial Septal Nuclei sub regions (MSN), (Figure S1), Thalamus (TH, mostly in lateral thalamus nuclei) and in the hippocampus (Hc) (predominantly in the CC, Figure S2). Damage consisted of extensive parenchymal inflammatory cells infiltration (lymphocytes, macrophages) in CC, MSN, TH brain regions and mild sporadic hemorrhages (detected only in CC area of the Hc). These findings were accompanied by extensive necrosis and degeneration.

The apparent safety of the vaccine when inoculated directly into the brain of C57BL/6J mice and, furthermore, to the sensitive IFNAR KO mice, confirms the safety of the vaccine with respect to the nervous system and strongly supports the overall safety profile of rVSV-ΔG-SARS-CoV-2-S vaccine.

## 4. Discussion

Beyond proving immunogenicity and efficacy, there is great importance to an in-depth characterization of the safety profile of a vaccine under development, in order to maintain public trust, especially in times of emergency. To this end, there is an advantage in choosing a vaccine platform which has already proven to be safe. In addition, the nonclinical safety profile should be carefully established in a variety of different models.

In light of the need for the development of vaccines for SARS-CoV-2 in a timely manner, we developed a replication competent recombinant VSV-ΔG-spike vaccine, in which the G gene which encodes the viral envelope glycoprotein of VSV was deleted from the viral genome and replaced with the S gene of SARS-CoV-2 [6], which mediates entry to cells expressing hACE2 receptor and is a major viral surface antigen to which neutralizing antibodies and T cell response are induced. The rVSV-ΔG-SARS-CoV-2-S vaccine utilizes the VSV backbone employed by other live vaccines including rVSV-EBOV which was approved by the FDA (under the trade name ERVEBO®), at the end of 2019, for the prevention of Zaire ebolavirus-caused disease in individuals 18 years of age and older [25]. The proven efficacy of the ERVEBO® vaccine to prevent Ebola infection in humans is remarkable and validates the VSV-ΔG backbone as a platform to produce effective and safe vaccines [26].

In order to evaluate the safety profile of rVSV-ΔG-SARS-CoV-2-S vaccine, we have completed a series of nonclinical safety, immunogenicity and efficacy (potency) studies in 4 animal species, mice, hamsters, rabbits and pigs, using multiple dose levels (up to 10^8^ PFU/animal). In addition, a repeat-dose GLP toxicity study was conducted in rabbits (out-with the scope of this article).

Particular attention was given to species selection, which was based on scientific, ethical and practical factors. According to the regulatory guidelines [7], a relevant animal species should develop an immune response similar to the expected human response after vaccination. Ideally, the animal species should also be sensitive to the pathogenic organism, and/or mimic the disease in humans. In line with these recommendations, the tested species presented increased levels of binding and neutralizing antibodies in response to vaccination, supporting their relevancy.

The first model to be established, the golden Syrian hamster (Mesocricetus auratus) model, was found to be susceptible to the COVID-19 disease and was used for evaluation of both the safety and efficacy of the vaccine [6]. In addition, safety was also evaluated in the K18 transgenic mouse model, which is considered a highly sensitive animal model for SARS-CoV-2. In these mice, human ACE-2 dependent replication of rVSV-ΔG-SARS-CoV-2-S is observed, followed by morbidity and mortality [27,28].

Interestingly, accumulated data places the hamster and the transgenic mouse model as the pivotal and indicative challenge models for SARS-CoV-2. While some other vaccine candidates have been tested in non-human primates (NHP), the different NHP strains tested show a variable response to the SARS-CoV-2 infection and thus results may be difficult to interpret with respect to determining the effectiveness and safety of the vaccine tested [28]. In addition, ethical considerations and limited availability of NHPs make this model less favorable and highly complex for use.

The rabbit, a non-rodent animal species, is considered suitably robust, sensitive and regulatory acceptable as an animal species for testing of vaccines, and was shown to develop an antibody signature response to SARS-CoV-2 [29]. In addition, the observed histopathological changes in our vaccinated rabbits (i.e. increased lymphocytic cellularity in the germinal centers of the spleen and regional lymph node) were considered to reflect a secondary change due to antigenic stimulation. Pigs, on the other hand, are more physiologically similar to humans than some other animal models in terms of body weight and metabolic rate. The pig is a valuable model for testing human safety and immunogenicity of vaccines [30], and may provide an indication of the type of immune response induced and is thus considered a valid nonrodent model for nonclinical safety testing. In addition, the pig model is considered to be an acceptable alternative to the dog model, which is commonly used as non-rodent toxicology model, or to non-human primates which are less accessible and less justified for ethical issues.

Altogether, the use of 4 different animal species, rodent and non-rodent, is important in building a reliable and comprehensive safety profile to support further clinical research.

Under the study conditions described here, rVSV-ΔG-SARS-CoV-2-S vaccine was found to be well tolerated, did not raise any safety signal of concern, up to the highest dose tested (10^8^ PFU) and evoked high levels of neutralizing antibodies against SARS-CoV-2, in all species tested.

No shedding of the viral vaccine was detected in urine or blood. There were no rVSV-SARS-CoV-2-S-related histopathological changes in hamsters and pigs, and the histopathological findings observed in rabbits are in line with what can be expected from an immunogenic vaccine, with immune stimulation (lymphoid hyperplasia in local lymph nodes and spleen, and mild local inflammation). No evidence for neurovirulence was found following intracranial injection of the vaccine (mimicking worst case conditions for maximum effect) in C57BL/6 immune competent mice or in IFNAR KO mice, deficient in the interferon alpha/beta receptor and hence highly sensitive to viral infections. Additionally, using the K18 hACE2 transgenic mouse model we demonstrated containment of the vaccine at the site of injection and its gradual clearance, with no distribution to the lungs. We also show that there is still viable vaccine virus up to 72 hours post vaccination in some of the animals – a time window which allows the formation and induction of a robust immune response. This is despite the fact that the receptor for the vaccine virus, namely hACE-2, is broadly expressed, including in the lungs, further supporting the safety profile of the vaccine candidate.

It is noteworthy that the emerging nonclinical safety profile of rVSV-ΔG-SARS-CoV-2-S vaccine is found to be very similar to that of the rVSV-EBOV [23,25,31,32]. A single dose of rVSV-EBOV did not lead to any biologically significant rVSV-ZEBOV-GP-related effects on clinical observations, mortality, body weights, body temperature, clinical chemistry and hematology in 2 animal models tested (mice and non-human primates). Similar treatment-related histopathological changes in injections site, iliac lymph nodes and spleen also indicate a similar mechanism of vaccine-induced antigenic stimulation.

One major potential concern associated with COVID-19 vaccines is the risk of Vaccine-Associated Enhanced Disease (VAED), including, but not limited to Vaccine Associated Enhanced Respiratory Disease (VAERD) which involves exacerbation of the respiratory viral infection’s clinical presentation, via antibody and complement-mediated mechanisms (namely Antibody-Dependent Enhancement, ADE) or via antibody-independent mechanisms which may involve cytokine release or cellular immune activation (recently reviewed [33]). As far as we know, to date, accumulating clinical data have shown no evidence of VAED/VAERD in any COVID-19 vaccine clinical trials, including those based on viral vectors. Nevertheless, VAED was previously noted in the development of Dengue virus, RSV [34], and measles vaccines [35]. In addition, pre-clinical data with other coronaviruses family members, SARS-CoV and Middle East respiratory syndrome (MERS) [36,37], also suggested signs of VAERD, when vaccinated animals were subsequently challenged with the wild type SARS‑CoV/MERS-CoV [38,39], and developed immunopathologic lung reactions that were attributed to a Th-2-biased immune response.

According to the recent FDA guidance document for development and licensure of COVID-19 vaccines [40], assessment of potential ERD in preclinical models should include an evaluation of neutralization versus total antibody responses, and a Th1/Th2 balance, whereby a vaccine showing a relatively high antibody titer of neutralizing antibodies, and an immune response that is primarily Th1, is considered at low risk for ERD. Accordingly, vaccination with rVSV-ΔG-SARS-COV-2-S vaccine led to high neutralizing antibody titers in all animal species tested, and a Th1-type T cell polarization was demonstrated in pigs, manifested by IFNγ, IL-2 and TNFα production but not Th2 cytokines (IL-4, IL-10) in spike-stimulated PBMCs. These results are consistent with what is generally known for other vaccines based on live viruses, specifically, for the VSV-∆G platform which was shown to elicit such a response [41].Furthermore, using two reliable SARS-CoV-2 challenge models, Syrian hamsters and K18 hACE2 transgenic mice (data not shown), we found that vaccination with rVSV-ΔG-SARS-COV-2-S vaccine protected the animals, and did not lead to disease enhancement, once the animals were subjected to a challenge with the wild type SARS-CoV-2 virus. Lungs of these animals were sustainably less damaged, and exhibited reduced viral load and improved in tissue to air ratio, as compared to infected, unvaccinated animals [6].

## 5. Conclusions

In conclusion, non-clinical data suggest that the rVSV-ΔG-SARS-COV-2-S vaccine is safe and immunogenic. These results supported the initiation of clinical trials (NCT04608305), currently in Phase 2.

## 6. Patents

Patent application for the described vaccine was filed by the Israel Institute for Biological Research.

## Acknowledgments

We acknowledge the IIBR administrative personnel for their commitment to the project. We thank Yossi Shlomovich, Noa Caspi and Ruchama Brody from IIBR animal husbandry; Yossi Lavie, Tal Levin-Harrus, Shkedia Kohen, Raphael Lioz and Boaz Nachshon from Envigo CRS (Israel) Ltd., Ness Ziona, Israel; Dr. Emmanuel Loeb from Patho-Logica Ltd; Dr. Tami Horovitz from Gsap, Israel for scientific editing of the manuscript; Prof. Eran Bacharach (Tel Aviv University) who kindly provided VSV-Indiana (WT-VSV) and the Bundeswehr Institute of Microbiology, Munich, Germany that kindly provided SARS-CoV-2 (GISAID accession EPI_ISL_406862).

