## Supplemental Figures for "Nonclinical Safety and Immunogenicity of an rVSV-ΔG-SARS-CoV-2-S vaccine in mice, hamsters, rabbits and pigs"

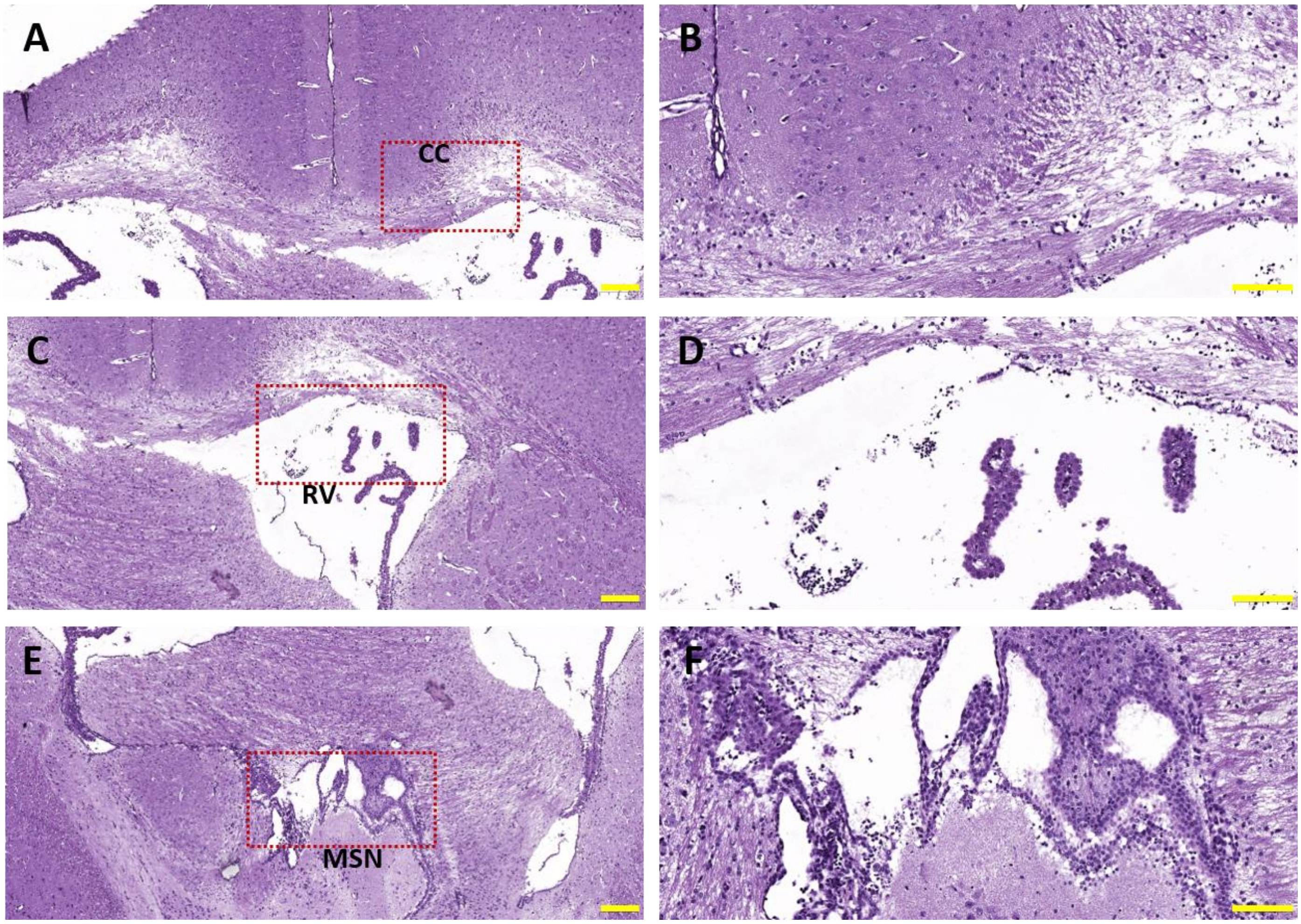


**Supplamnatry Figure 1: Histopathology of C57/BL6 mouse brain striatum after injection VSV-WT**

WT C57/BL6 striatum brain area: (A-B) Corpus Callosum, CC; (C-D) Right Ventricle, RV; (E-F) Medial Septal Nuclei, MSN. H&E staining. Scalebar=200µm, Magnification: X5 (A, C, E); Scale bar=100µm, Magnification X15 (B, D, F).

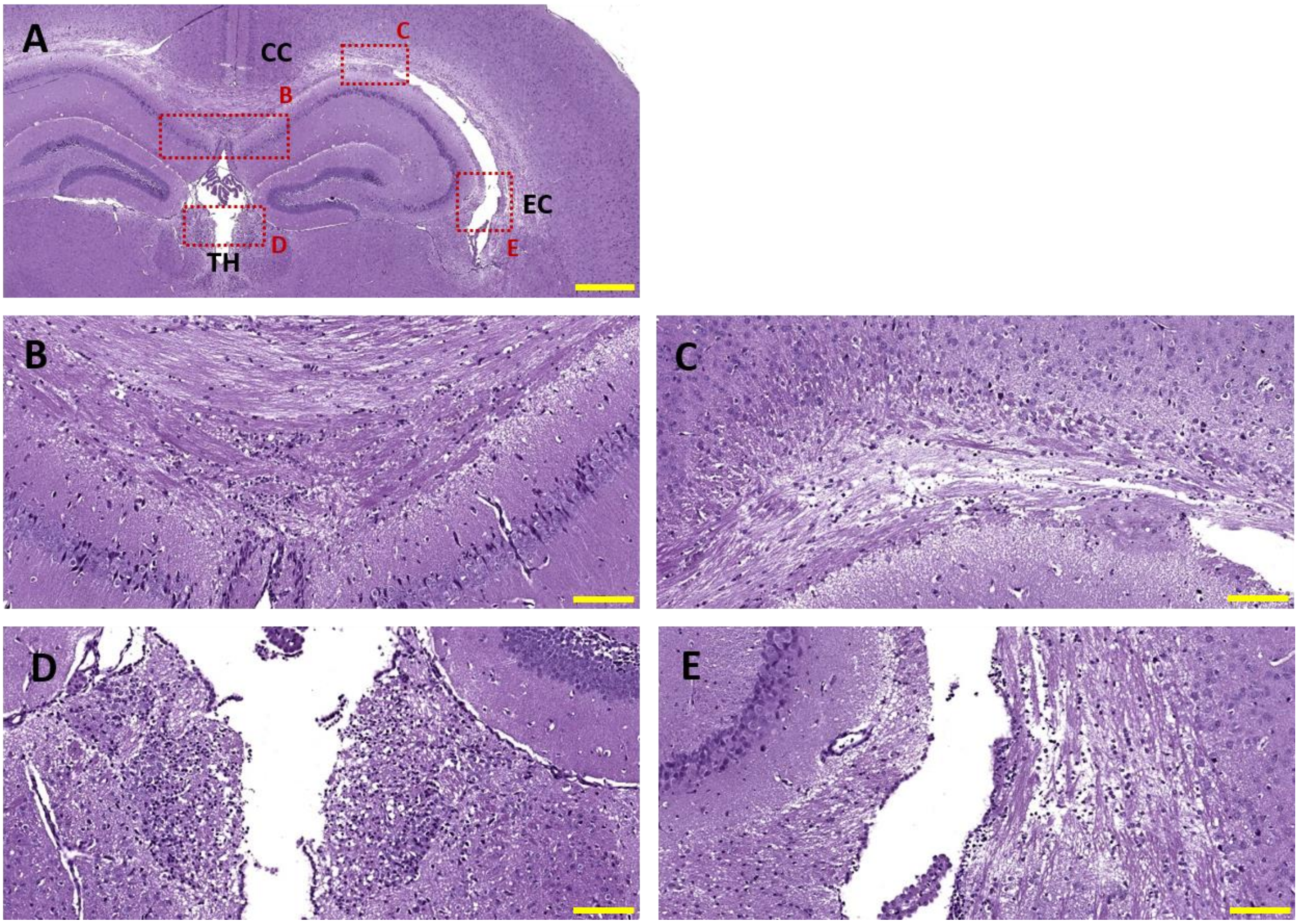


**Supplementary Figure 2: Histopathology of C57/BL6 mouse brain thalamus and hippocampus after injection with VSV-WT**

WT C57/BL6 Hc and TH brain areas: (A) Hc, in squares: damaged areas. (B-C) Corpus Callosum, CC; (D) TH - lateral; (E) External capsule, EC. H&E staining: (A): Scale bar=500µm, Magnification: X3; (B-E): Scale bar=100µm, Magnification X15.
